# CRHR1-mediated Akt activation involves endocytosis and soluble adenylyl cyclase activity

**DOI:** 10.1101/2022.08.04.502800

**Authors:** Paula A. dos Santos Claro, Natalia G. Armando, Alejandra Attorresi, Karen E. Lindl, Micaela Silbermins, Carolina Inda, Susana Silberstein

## Abstract

Corticotropin-releasing hormone receptor 1 (CRHR1) signaling initiates at the cell membrane and continues after receptor internalization. A sustained cAMP generation after ligand-activated CRHR1 endocytosis supported by the soluble adenylyl cyclase (sAC) is required for the G protein-independent phase of ERK1/2 phosphorylation in neuronal cells. Here, we report that Akt is activated by CRHR1 stimulation in a mechanism dependent on the endosomal cAMP production by sAC activity. Moreover, AktS473 phosphorylation profile was distinct to that of phospho-ERK1/2 revealing a crosstalk between these pathways. We found that the CRHR1 activation of PI3K/Akt pathway is required for a full ERK1/2 activation but negatively regulates the cAMP response and CREB phosphorylation downstream CRHR1 stimulation. Immunofluorescence colocalization analysis revealed that activated CRHR1 transits to early (Rab5) and recycling (Rab11) endosomes. Additionally, CRHR1 colocalized with two Rab5 effectors, APPL1 and EEA1. shRNA mediated knockdown uncovered a role of these proteins in CRHR1 signaling. APPL1 silencing led to a decrease in pAktS473 levels without affecting ERK1/2 activation, whereas EEA1 depletion had the opposite effect, suggesting that CRHR1 intracellular signaling diversity may be achieved through different signaling platforms.

## INTRODUCTION

Corticotropin-releasing hormone (CRH) and the CRH-related peptide urocortin1 are high affinity ligands that activate the class B/secretin-like GPCR corticotropin-releasing hormone receptor 1 (CRHR1), that signals mainly by Gs coupling, leading to cyclic AMP (cAMP) increase and activation of multiple signaling pathways (Inda et al., 2017a). Ligand-activated CRHR1 continues to signal even after internalization in neuronal contexts (Inda et al., 2017b), operating by different signaling effectors, some widely known but also other atypical (Inda et al., 2016). Thus, activated CRHR1 signaling properties are in line with recent evidence supporting the paradigm of GPCR signaling from different cellular locations, which lead to the notion of temporal and spatial encoding of cellular responses.

After ligand-dependent internalization, receptors are transported through the endosomal network. In general, cargoes such as cell surface receptors, are internalized to Rab5 early endosomes from which they can either be recycled back to the plasma membrane in Rab4/Rab11 recycling endosomes or sent to degradation in Rab7/Rab9 late endosomes (von Zastrow and Sorkin, 2021). Rab proteins confer functional and structural identity to the endocytic network, mainly by recruiting specific effector proteins to a particular intracellular compartment (Sigismund et al., 2012).

CRH plays a central role in the regulation and integration of neuroendocrine, autonomic, and behavioral adaptive responses to stress. Hypothalamic CRH-secreting neurons drive both basal and stress-induced activation of the hypothalamic-pituitary-adrenal (HPA) axis. In addition, CRH is widely distributed in the brain, where it acts as a neuromodulator, integrating a complex system that regulates the behavioral stress response. Dysregulation of CRH action through CRHR1 is crucial in the pathogenesis of psychiatric conditions (i.e. depression, anxiety, addictions), neuroendocrinological alterations, and during the onset and development of neurodegenerative diseases (Deussing and Chen, 2018).

Using mouse hippocampal HT22 cells stably expressing CRHR1 (HT22-CRHR1 cells) as a cellular model, we have been able to define fundamental molecular mechanisms underlying CRHR1 signaling in a neuronal context. We described that the temporal pattern of ERK1/2 phosphorylation in response to CRH is biphasic, with an early acute peak 3-6 min after stimulation-dependent on B-Raf and protein kinase A (PKA)- and a second sustained phase that remained active for at least 60 min after CRH addition, dependent on CRHR1 internalization and β-arrestin2 (Bonfiglio et al., 2013). Importantly, in the hippocampal context as well as in corticotrophs, ERK1/2 activation was dependent on cAMP levels in response to CRH (Bonfiglio et al., 2013; Kovalovsky et al., 2002).

Our recent work demonstrated that CRHR1 activates soluble adenylyl cyclase (sAC) besides transmembrane adenylyl cyclases (tmACs) in neuronal and endocrine cells (Inda et al., 2017b; Inda et al., 2016). Only sAC activity is essential for internalization-dependent cAMP production and sustained ERK1/2 activation responses in HT22-CRHR1 cells, revealing a functional association between sAC-generated cAMP and endosome-based GPCR signaling. We provided the first evidence of sAC being involved in an endocytosis dependent cAMP response, supporting the current model of GPCR signaling in which the cAMP response does not exclusively occur at the plasma membrane, and introducing the notion of sAC as an alternative source of cAMP. Furthermore, cAMP-dependent CRHR1 signaling response was found sustained in hippocampal neuronal cells, but transient and acute in corticotrophs, highlighting that the CRH system signaling pathways are specific of cellular context both in terms of effectors and signal duration.

The study of Akt kinase signaling is in a remarkable expansion due to its relevance in numerous pathologies such as cancer and neurological disorders. Akt activity constitutes an emblematic example of the transformation of extracellular information in essential cell decisions on metabolism, protein synthesis, growth and survival (Manning and Toker, 2017). In the central nervous system, PI3K/Akt pathways play a role in neuron development, morphology, establishment and conservation of neural circuits (Hou and Klann, 2004). In the hippocampus, Akt activity and its substrates play a key role in synaptic plasticity (Levenga et al., 2017) whose dysregulation impacts on memory and cognitive impairments which are frequently associated to psychiatric and neurological illness (Manning and Toker, 2017). Akt substrates, such as GSK3, were found to mediate the action of psychoactive drugs such as antidepressants, antipsychotics, and other mood stabilizer compounds (Matsuda et al., 2019).

In the present study, we aimed to characterize PI3K/Akt signaling in the hippocampal neuronal context of CRHR1 action. The temporal profile of Akt phosphorylation on S473 is consistent with an activation stimulated by internalized CRHR1. Consistently, the cAMP pool involved in Akt activation is provided by sAC, the critical source responsible for the internalization-dependent production of this second messenger for CRHR1. Moreover, blocking endocytosis impairs Akt activation. The PI3K/Akt pathway is required for complete ERK1/2 activation by CRHR1 stimulation in HT22-CRHR1 cells. Interestingly, by measuring the cAMP response triggered by activated CRHR1 using a Förster resonance energy transfer (FRET)–based biosensor, we found that pharmacological inhibition of PI3K and Akt increases cAMP levels. Fluorescent immunocytochemical analysis allowed the detection of CRHR1 spatially coincident with endogenously expressed Rab proteins characteristic of early and recycling endosomes, and Rab5 effectors APPL1 and EEA1. Moreover, APPL1 and EEA1 knockdown differentially affects Akt and ERK1/2 activation, suggesting that CRHR1 signaling from early endosomal compartments involves several specific signaling platforms.

## RESULTS

### CRHR1 activation induces Akt phosphorylation and activity in HT22-CRHR1 cells

First, we explored whether ligand-stimulated CRHR1 was able to trigger Akt activation in HT22-CRHR1 cells by immunodetection of phosphorylated Akt at S473 site (pAktS473) and total Akt in cell lysates. Stimulation with 10nM CRH or UCN1 triggered Akt phosphorylation (Fig. 1A). pAktS473 levels were robustly evidenced 20 minutes after CRHR1 activation and remained constant after 60 minutes of ligand incubation. High affinity ligands for CRHR1 (CRH and UCN1) elicited Akt activation, whereas specific ligands for CRHR2 (UCN2 and UCN3) did not (Fig. S1A), confirming that the response depends on activated CRHR1, and being consistent with the lack of CRHR2 expression in this cell line.

**Figure 1:**
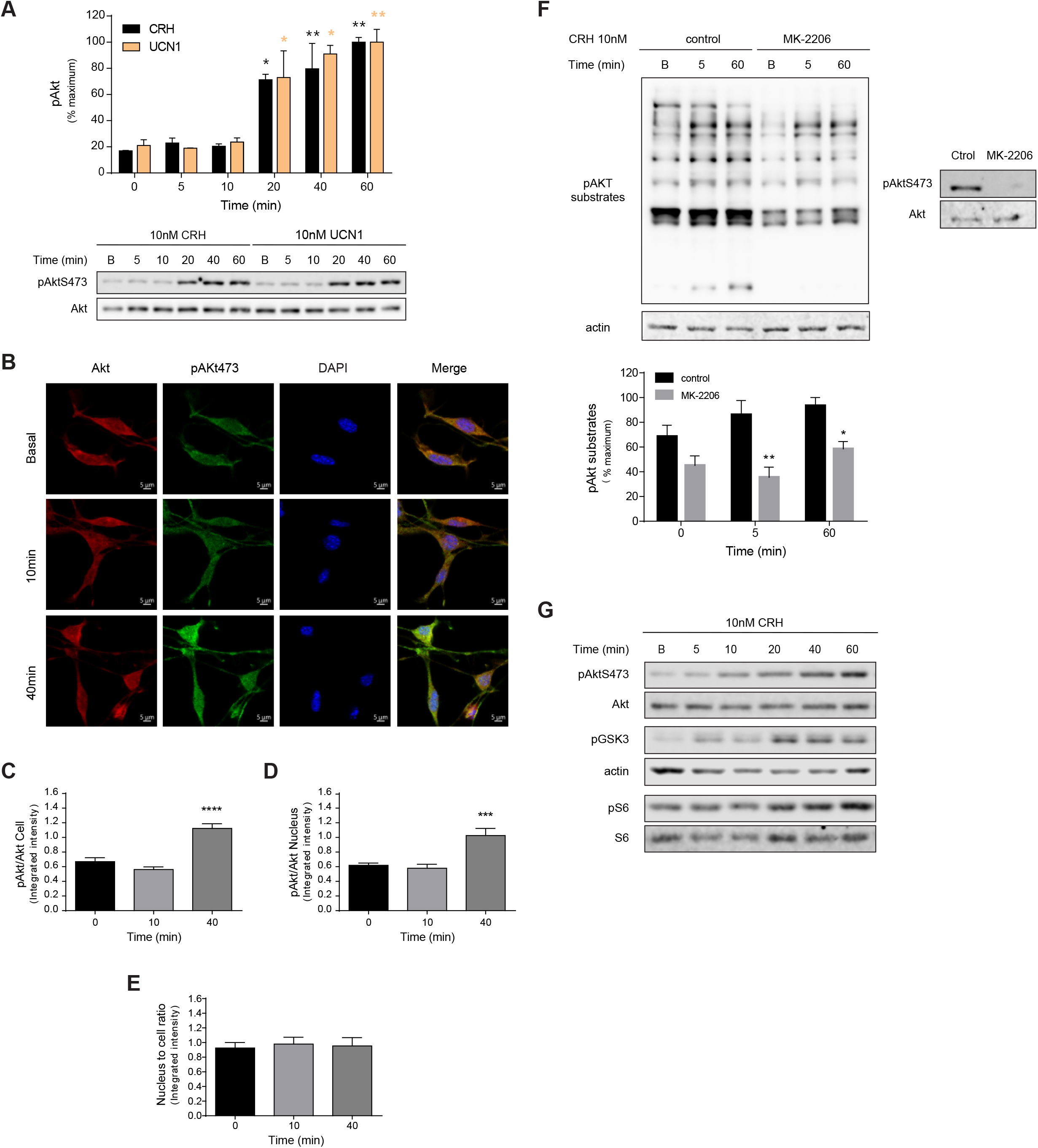
Activated CRHR1 promotes Akt phosphorylation and activity in HT22-CRHR1 cells. A) pAktS473 and total Akt were measured in HT22-CRHR1 cells stimulated with 10nM CRH or 10 nM UCN1 for the indicated time points. Signals were quantified by densitometry and each value of pAkt was normalized to total Akt. Results are expressed as percentage and relative to the mean of maximum pAkt (mean ± s.e.m, n=3). Two-way analysis of variance followed by Bonferroni’s test: *, p < 0.05; **, p < 0.01 with respect to basal of each condition. Representative immunoblots are shown. B) Akt and pAktS473 subcellular distribution were analyzed by immunofluorescence in cells stimulated with 10 nM CRH for the indicated time points using specific antibodies against pAktS473 (green) and total Akt (red). Nuclei were stained using DAPI. Representative images obtained for each condition are shown. Bars, 5μM. C-E) pAktS473 and total Akt fluorescent signals were quantified in cell (C) and nuclei (D) using *CellProfiler* software and each value of pAkt was normalized to total Akt. Nuclei to cell ratio was calculated (E). Data: mean ± s.e.m, n =3, 30-40 cells per condition. One-way analysis of variance followed by Tukey’s test: ***, p < 0,001; ****, p < 0,0001; with respect to basal. F) pAkt substrates phosphorylation was immunodetected in cells preincubated with vehicle (control) or Akt-specific inhibitor (10μM MK-2206). pAkt and total Akt were detected as a control of inhibitor’s efficiency. Actin was detected as a loading control. Results are expressed as the percentage of maximum response. (mean ± s.e.m, n = 3). Two-way analysis of variance followed by Bonferroni’s test: *, p < 0.05; **, p < 0.01 with respect to control. G) pAktS473, pGSK3, pS6 and their respective total proteins were determined by immunoblotting in HT22-CRHR1 cells stimulated with 10nM CRH. Representative immunoblots are shown.

We examined the spatial distribution of Akt and pAktS473 by indirect immunofluorescence in HT22-CRHR1 cells (Fig. 1B). An increase of the pAktS473 signal after CRHR1 stimulation was evidenced with a similar temporal profile to the one detected in cell lysates. Given that Akt and pAkt signals were distributed throughout the cell, we performed nuclear segmentation using DAPI to quantify Akt activation in the whole cell or in the nucleus. pAktS473 fluorescent intensity relative to total Akt increased by about 50% in response to CRH stimulation in both compartments (Fig. 1C,D). Nuclei to cell ratio remained constant, indicating that this rise in pAkt levels occurs evenly throughout the cell (Fig. 1E).

Consistent with the activation of Akt kinase function downstream activated CRHR1, immunoblots of CRH-stimulated HT22-CRHR1 cells lysates showed elevated levels of several Akt phospho-substrates, which was prevented in the presence of the Akt-specific inhibitor MK-2206 (Fig. 1F). Additionally, we examined the time course of the phosphorylation profile of GSK3 and S6, two proteins known to be activated by pAkt. Phosphorylation levels of these well-known Akt substrates in response to CRH temporally correlated with the increase of pAkt (Fig. 1G). Thus, Akt activity is triggered downstream stimulated CRHR1 in HT22-CRHR1 cells.

### CRHR1-induced Akt activation requires cAMP: a critical role of sAC

Since we have previously demonstrated that CRH stimulation results in a rapid increase of intracellular cAMP levels in HT22-CRHR1 cells, as well as in primary neurons (Inda et al., 2017b; Inda et al., 2016), we assessed if Akt activation was dependent on the cAMP response in these cells.

We determined that CRH and UCN1-stimuli capable of triggering Akt activation-elicited a rapid and sustained cAMP response as evidenced by FRET changes in HT22-CRHR1 cells stably expressing the FRET-based biosensor Epac-S^H187^ (HT22-CRHR1-Epac-S^H187^) (Inda et al., 2017b; Inda et al., 2016), whereas specific ligands for CRHR2 (UCN2 and UCN3) did not trigger a cAMP response (Fig. S1B). Treatment with the cell membrane-permeable cAMP analogue 8-CPT-cAMP or with the receptors-independent activator of tmACs forskolin induced Akt phosphorylation in these cells (Fig. 2A). Notably, the temporal pattern of Akt activation obtained by increasing intracellular cAMP levels was similar to that of CRH-stimulated CRHR1. These results suggest that Akt activation is dependent on CRH increased cAMP levels in HT22-CRHR1 cells.

**Figure 2:**
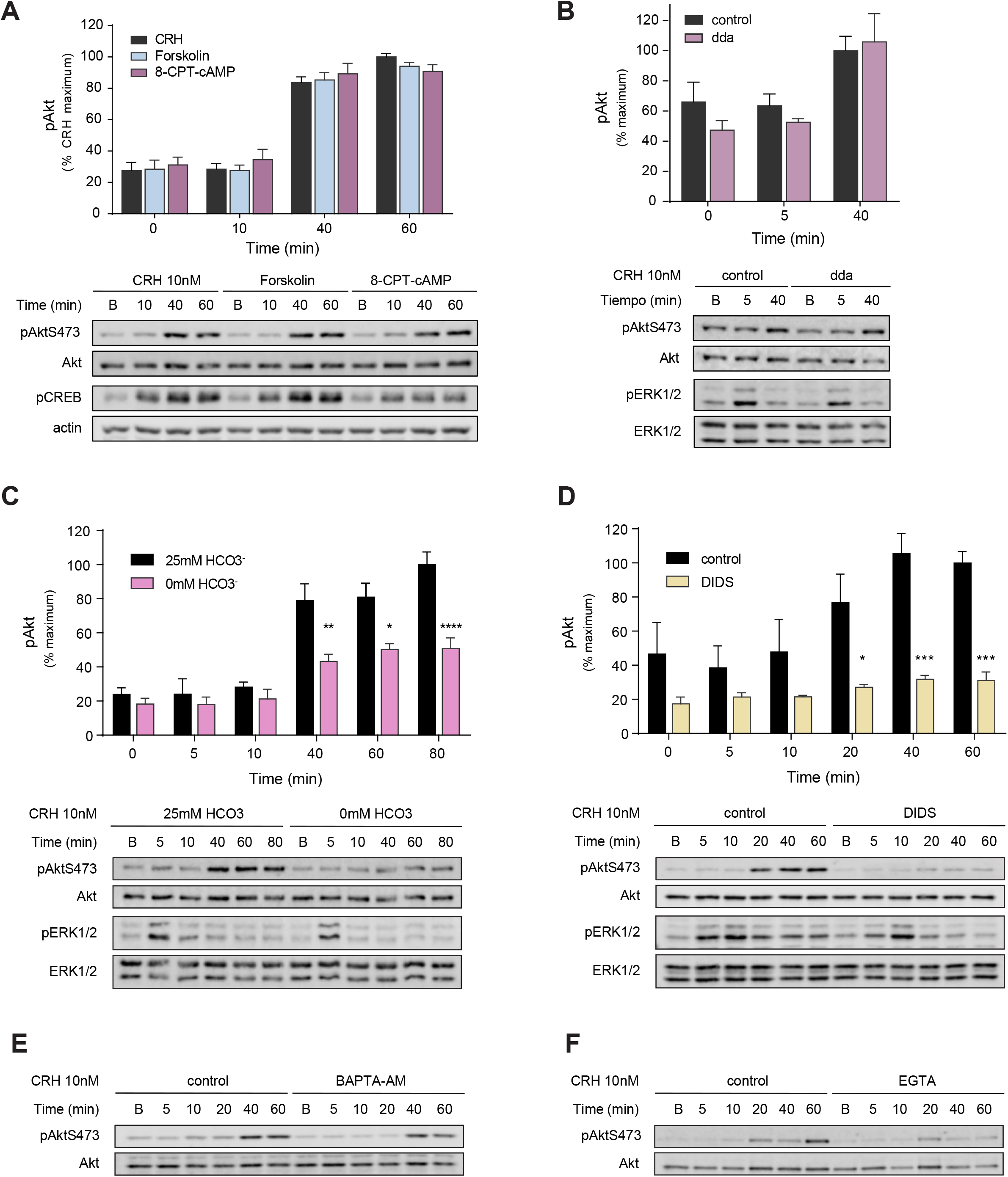
Akt phosphorylation is dependent on sAC-generated cAMP. A) HT22-CRHR1 cells were stimulated with 10 nM CRH, or treated with 50 μM forskolin or 50 μM 8-CPT-cAMP for the indicated time points. Phosphorylated and total proteins were determined by immunoblotting. Signals were quantified by densitometry and each value of pAkt was normalized to total Akt. pCREB was detected as a control of stimuli efficiency and actin as a loading control. Results are expressed as percentage and relative to the mean of maximum pAkt with CRH stimulation (mean ± s.e.m, n=4). Representative immunoblots are shown. (B-F) HT22-CRHR1 cells stimulated with 10 nM CRH for the indicated time points. Phosphorylated and total proteins were determined by immunoblotting. Signals were quantified by densitometry and each value of pAkt was normalized to total Akt. pERK1/2 and total ERK1/2 are presented as a treatment control. Results are expressed as percentage and relative to the mean of maximum pAkt in control condition. Representative immunoblots are shown. B) Cells were pretreated with vehicle (DMSO) or tmACs inhibitor (10 μM 2’-5’ dda). Data: mean ± s.e.m, n=3. C) Cells were stimulated in HCO3-free or 25 mM HCO3-DMEM. Data: mean ± s.e.m, n=3. Two-way analysis of variance followed by Bonferroni’s test: *, p < 0.05; **, p < 0.01; ****, p < 0.0001 with respect to control. D) Cells were pretreated with vehicle (control) or DIDS (450 μM). Data: mean ± s.e.m, n=3. Two-way analysis of variance followed by Bonferroni’s test: *, p < 0.05; ***, p < 0.001; with respect to control. E) Cells were pretreated with vehicle (DMSO) or calcium permeable chelator BAPTA (5 μM). F) Cells were pretreated with vehicle (control) or extracellular calcium chelator EGTA (500 μM).

We have reported that the CRHR1-triggered cAMP response does not exclusively depend on classic transmembrane adenylyl cyclases (tmACs) but also involves the soluble enzyme sAC as a second source of cAMP (Inda et al., 2016). Therefore, the dependence of Akt phosphorylation on different cAMP sources was investigated (Fig. 2 B-F). First, we assayed the effect of blocking cAMP generation from tmACs on Akt and ERK1/2 activation by CRH. As previously reported, specific inhibition of tmACs with ddA attenuated the CRH-induced early (5 min) ERK1/2 activation phase (Bonfiglio et al., 2013), but had no effect on Akt phosphorylation in response to CRH (Fig. 2B).

Next, we examined the effect of the sAC-specific modulators bicarbonate (Kleinboelting et al., 2014; Kleinboelting et al., 2016; Maitan et al., 2021; Steegborn et al., 2005) and calcium (Jaiswal and Conti, 2003; Litvin et al., 2003) on the Akt response to CRH stimulation. HT22-CRHR1 cells stimulated in bicarbonate-free medium exhibited an attenuated ERK1/2 activation (Inda et al., 2016). Remarkably, in absence of bicarbonate, Akt activation was severely impaired (Fig. 2C) as well as Akt substrates phosphorylation (Fig. S2). Pharmacologic treatment with DIDS inhibitor of bicarbonate transporters, as well as sAC activity by competition with bicarbonate binding (Kleinboelting et al., 2014), caused an almost complete blockage of Akt activation as well (Fig. 2D). Consistent with the requirement of calcium to activate sAC, Akt activation downstream stimulated CRHR1 was diminished by preincubating cells with the calcium chelators EGTA and BAPTA-AM (Fig. 2E,F). Thus, we conclude that the phosphorylation of Akt 473 site by CRH is dependent on increased cAMP levels by CRHR1 activation of bicarbonate- and calcium-dependent sAC.

### CRHR1-induced Akt activation modulates the ERK1/2 response

Our results are consistent with a mechanism of Akt activation downstream CRHR1-triggered cAMP in which sAC is the main source of cAMP involved. Given that Akt activation was evidenced at late time points of CRHR1 stimulation, we compared the temporal patterns of Akt and ERK1/2 kinases phosphorylation. A distinct profile of Akt S473 phosphorylation compared to phospho-ERK1/2 was evident after CRH stimulation in HT22-CRHR1 cells (Fig. 3A). As we previously reported, CRH induced ERK1/2 activation in a biphasic manner (Bonfiglio et al., 2013). Akt phosphorylation was not detected at early time points when pERK1/2 reaches maximal levels but increased during the late time points in which the sustained phase of ERK1/2 activation is evident, reaching its maximum level by 60 min of CRH stimulation (Fig. 3A). Different temporal profiles of Akt and ERK1/2 activation were also observed using PDGF as a stimulus, which rapidly activated both kinases activities (Fig. S3).

**Figure 3:**
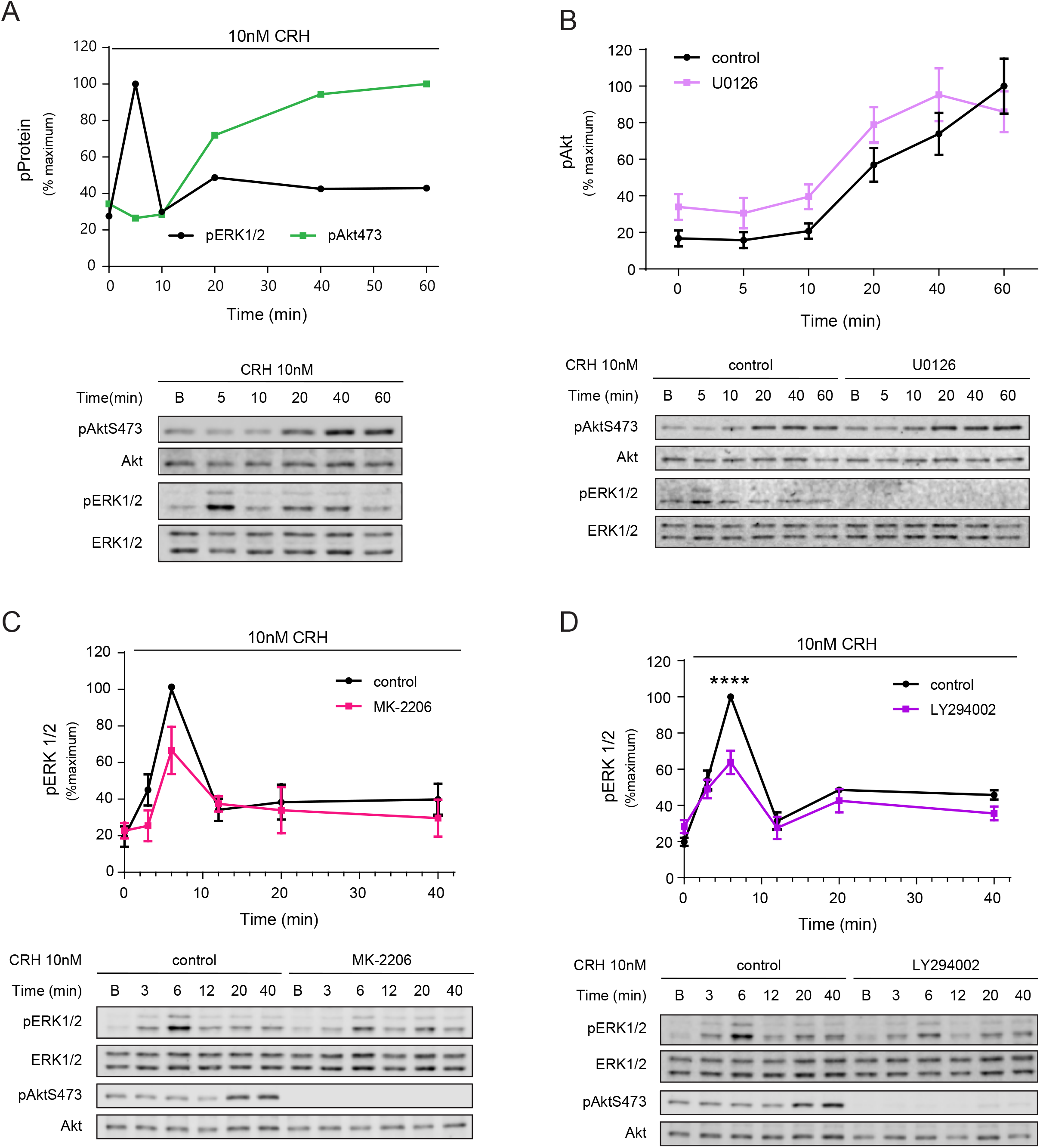
Akt and ERK1/2 crosstalk in CRH-activated HT22-CRHR1 cells. A) HT22-CRHR1 cells were stimulated with 10 nM CRH for the indicated time points. Phosphorylated and total proteins were determined by immunoblotting. Signals were quantified by densitometry and each value of pAkt, or pERK1/2 was normalized to total Akt or ERK1/2, respectively. Representative immunoblots are shown. Results are expressed as percentage of maximum pAkt or pERK1/2. B-E) Cells were preincubated with vehicle (control) or (B) MEK-specific (10μM U0126) or (C) PI3K-specific (20 µM LY294002) or (D) Akt-specific (10μM MK-2206) inhibitors, before stimulation with 10 nM CRH. B) pERK1/2 and total ERK1/2 were detected as a control of inhibitor’s eficiency. Results are expressed as percentage and relative to the mean of maximum pAkt in control condition. Data: mean ± s.e.m, n=4. C, D) pAktS473 and total Akt were detected as a control of inhibitor’s efficiency. Results are expressed as the percentage of maximum response in control conditions. Data: mean ± s.e.m, n=4. Two-way analysis of variance followed by Bonferroni’s test: ****, p < 0.0001 with respect to control. Representative immunoblots are shown.

Akt and ERK1/2 signaling pathways interact to shape multiple cell responses (Mendoza et al., 2011). These signaling cascades often influence each other both negatively and positively through complex crosstalk mechanisms. The different profiles of Akt and ERK1/2 phosphorylation in response to CRHR1 activation may be reflecting a cross-regulation between both pathways. To investigate this possibility, we tested the effect of blocking each kinase activation by pre-treating cells with specific inhibitors before CRH stimulation.

The MEK1/2-specific inhibitor U0126 used at a concentration that completely blocked ERK1/2 phosphorylation, slightly increased pAktS473 levels triggered by CRH, indicating that the ERK1/2 pathway does not significantly affect Akt activation (Fig. 3B). Conversely, pharmacological blockage of Akt activation with specific inhibitor MK-2206, caused a diminished pERK1/2 response (Fig. 3 C).

Akt can be phosphorylated and activated by PI3K-dependent and -independent mechanisms (Clement et al., 2018). To determine PI3K involvement, we preincubated cells with the PI3K specific inhibitor LY294002. Treatment with LY294002 blocked Akt phosphorylation at the 473 site and strongly diminished ERK1/2 phosphorylation levels at both early and late phases (Fig. 3D). Therefore, PI3K/Akt activation is required for a full ERK1/2 activation by CRHR1 stimulation in HT22-CRHR1 cells.

Overexpression of wild-type Akt1 or a constitutively active Akt1 (E17K) form (Carpten et al., 2007) did not change pERK1/2 levels induced by CRH in HT22-CRHR1 cells, although very high levels of Akt and pAktS473 were immunodetected in lysates from cells transfected with these constructs (Fig. S4). Given the negative effect of PI3K/Akt inhibition on ERK1/2 activation (Fig. 3C,D), an increase in pERK1/2 levels could be predicted in these experiments. However, Akt1 overexpression and activation in the cell may be taking place at cellular locations different from those of ERK1/2, making the kinase unable to modulate ERK1/2 activity. Alternatively, an intermediate component linking Akt to ERK1/2 may be limiting, or another Akt protein (Akt2 or Akt3) can be the specific mediator of pAkt effect on ERK1/2 in CRHR1 signaling.

### CRHR1-induced PI3K/Akt activation modulates the cAMP response

Given that CRHR1 stimulation triggered a cAMP-dependent Akt activation through sAC (Fig. 2) and that the PI3K/Akt pathway was required for a full ERK1/2 response (Fig. 3C,D), we investigated the effect of the PI3K/Akt pathway activation on the cAMP response to CRH in HT22-CRHR1-Epac-S^H187^ cells (Inda et al., 2016). The CRH-triggered cAMP production assessed at single-cell level in real time, without phosphodiesterase inhibitors resulted in a rapid increase of intracellular cAMP levels after CRH application that remained elevated during the time course (Inda et al., 2016). When the PI3K-specific inhibitor LY294002 was added during the time course, FRET responses were increased evidencing higher levels of intracellular cAMP without changes in the temporal profile of the response (Fig. 4A). The addition of the Akt-specific inhibitor MK-2206 also augmented cAMP levels, although to a minor extent (Fig. 4B), most probably due to the non-optimal conditions used for FRET measurement to avoid interference caused by the intrinsic fluorescence of MK-2206 (see Materials and Methods).

**Figure 4:**
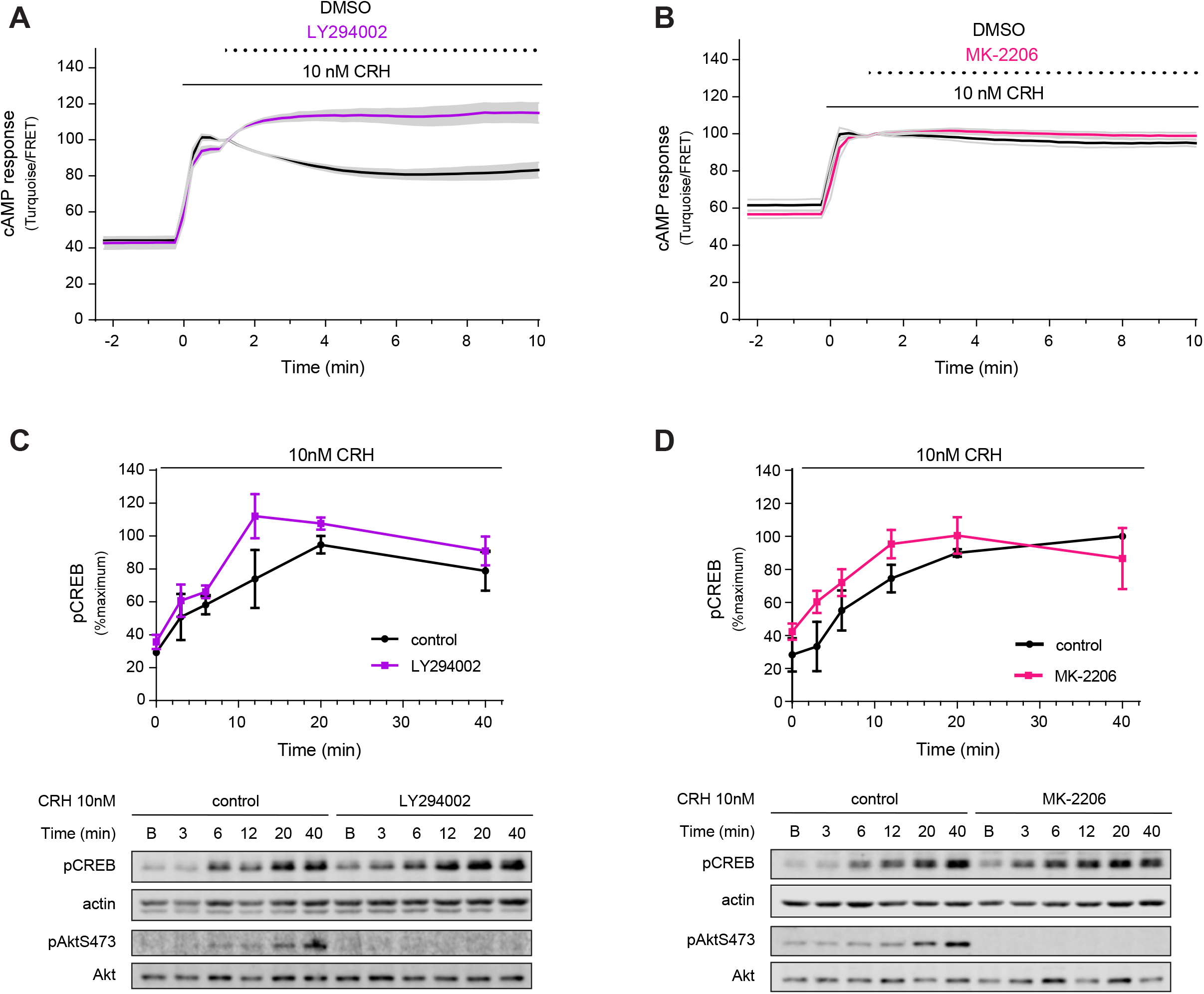
Inhibition of PI3K/Akt pathway increases CRH-mediated cAMP response and CREB phosphorylation. A,B) Time course of FRET changes measured in HT22-CRHR1-Epac-S^H187^ cells. Cells were stimulated at time 0 with 10nM CRH and vehicle (DMSO) or (A) PI3K-specific (20µM LY294002) or (B) Akt-specific (10μM MK-2206) inhibitors were added 1.5 minutes later (mean ± s.e.m, n=3). FRET conditions were established depending on the inhibitor tested (see Materials and Methods). C-D) HT22-CRHR1 cells were stimulated with 10nM CRH for the indicated time points and preincubated in the presence of vehicle (control) or (C) PI3K-specific (20µM LY294002) or (D) Akt-specific (10µM MK2206) inhibitors. Phosphorylated and total proteins were determined by immunoblotting. Signals were quantified by densitometry and each value of pCREB was normalized to actin. pAktS473 and total Akt are included as a control of inhibitor’s efficiency. Results are expressed as percentage and relative to the mean of maximum pCREB in control condition (mean ± s.e.m, n=4).

The cAMP-response element binding protein CREB, a crucial regulator of neuronal function, is the archetypal transcription factor activated by cAMP, which is a main effector of CRHR1 signaling (Inda et al., 2017b). Consistent with enhanced cAMP levels caused by inhibiting PI3/Akt signaling, CRH-stimulated HT22-CRHR1 cells showed increased phosphorylation of CREB at S133 (Fig. 4C,D).

Together, these results suggest that PI3K/Akt pathways modulate the cAMP response elicited by the activated CRHR1. However, the cAMP increase caused by PI3K/Akt inhibitors does not result in an increase in pERK1/2 levels, which are dependent on cAMP (Bonfiglio et al., 2013; Inda et al., 2016) and Akt for full activation (Fig. 3 C,D). A possible explanation for this paradox is that cAMP generation resulting from PI3K/Akt inhibition constitutes a spatially restricted pool unable to mediate ERK1/2 activation. This is highly plausible in the context of cAMP compartmentalization where cAMP domains are able to function as independent signaling units within a cell (Anton et al., 2022).

### Regulators of Akt activation by CRHR1 in HT22-CRHR1 cells

Pharmacologic or genetic inhibition of classic cAMP effectors, PKA and EPACs, had no effect on pAktS473 levels in response to CRHR1 (Fig. S5). We then investigated the role of Src kinase, an important upstream regulator of Akt and ERK1/2 pathway activation downstream GPCR signaling (Roskoski, 2015). We first tested the effect of Src family kinase (SFK) inhibitors on ERK1/2 and Akt activation stimulated by CRH in HT22-CRHR1 cells. Preincubation with Src inhibitor SU6656 caused a diminished Akt activation and a remarkably attenuated ERK1/2 response to CRH (Fig. 5A,B). Similar results were obtained with other two established Src inhibitors, Saracatinib and Dasatinib (Fig. 5C), suggesting that Src is involved in Akt and ERK1/2 activation by CRHR1. Other kinases, different from Src, are known to be affected by these small molecule inhibitors (Blake et al., 2000; Roskoski, 2015). This fact could account for the stronger effect of pharmacologic inhibition on pERK1/2 levels. To further analyze the involvement of Src on Akt response to CRH, cells were transfected with expression vectors of an active (SrcYF) or a dominant-negative (SrcYF-KM) form of the Src kinase (Servitja et al., 2003). Expression of SrcYF increased pAktS473 levels whereas SrcYF-KM blocked Akt activation in CRH-stimulated HT22-CRHR1 cells (Fig. 5D). On the other hand, ERK1/2 activation was not affected by these treatments.

**Figure 5:**
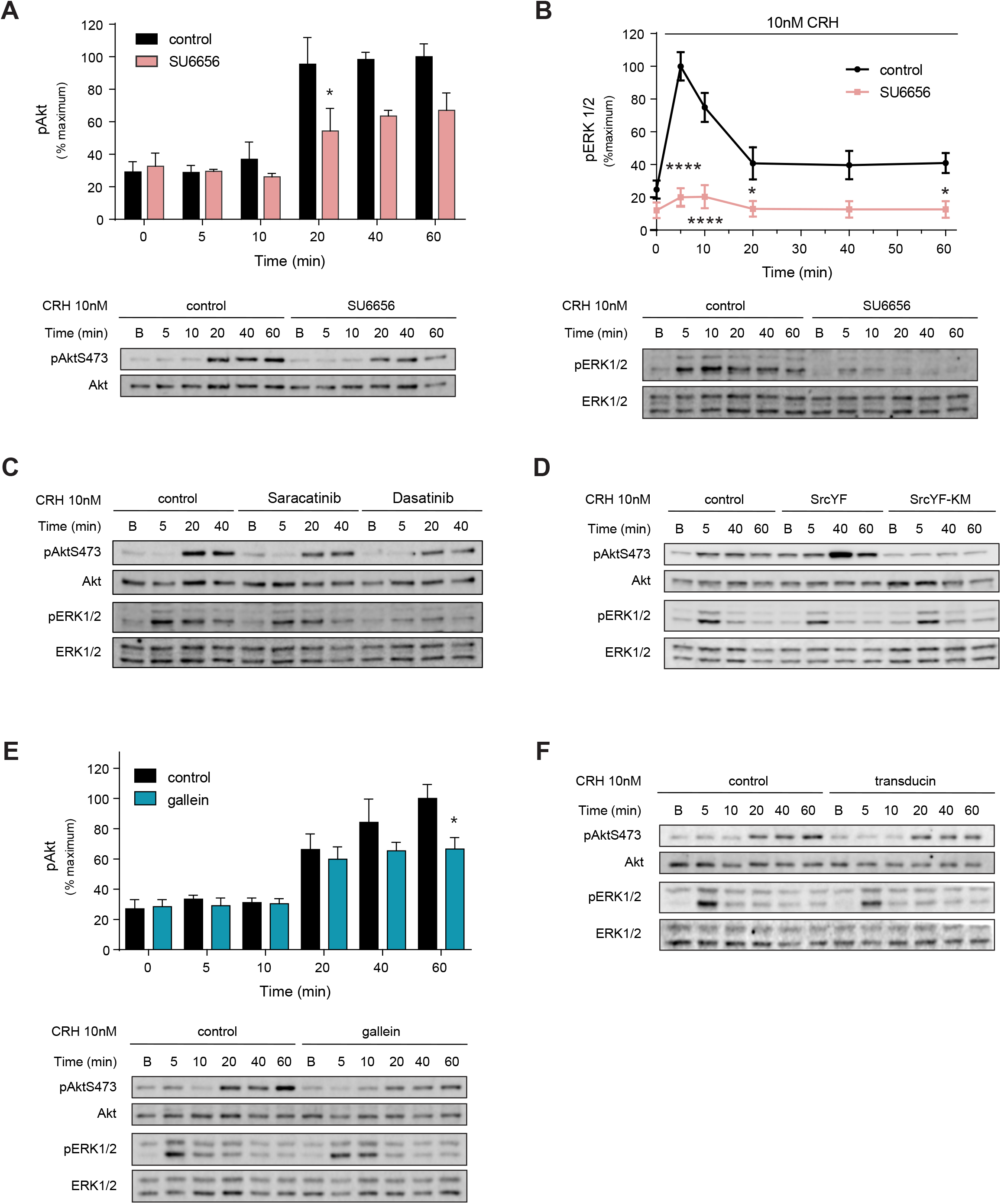
Src kinase and the Gβγ complex mediate Akt activation in response to CRH. A,B) HT22-CRHR1 cells were preincubated in the presence of vehicle (control) or SFK family inhibitor SU6656 3μM before stimulation with 10 nM CRH for the indicated time points. Phosphorylated and total proteins were determined by immunoblotting. Signals were quantified by densitometry and each value of pAkt (A), or pERK1/2 (B) was normalized to total Akt or ERK1/2, respectively. Representative immunoblots are shown. C,E) Cells were preincubated in the presence of vehicle (control) or SFK inhibitors 100 nM Saracatinib and Dasatinib (C) or Gβγ-specific inhibitor 10µM gallein (E) before stimulation with 10 nM CRH for the indicated time points. Phosphorylated and total proteins were determined by immunoblotting. E) Results are expressed as percentage and relative to the mean of maximum response in control condition (mean ± s.e.m, n = 3). Two-way analysis of variance followed by Bonferroni’s test: *, p < 0,05; ****, p < 0,0001 with respect to control. D, F) Cells were transfected with expression plasmids Src mutants, pCEFL-SrcYF or pCEFL-SrcYF-KM (D) or Gβγ scavenger pCDNA3-Gα transducin (F) 48hs before stimulation. Phosphorylated and total proteins were determined by immunoblotting.

Gβγ is often associated to Src activation downstream of GPCRs. In COS7 and HEK293 cells overexpressing CRHR1, CRH was shown to activate PI3K/Akt pathways in a Gβγ-dependent mechanism (Parra-Mercado et al., 2019; Punn et al., 2006). In order to evaluate the potential implication of Gβγ on Akt and ERK1/2 activation downstream stimulated CRHR1 in HT22-CRHR1 cells, we tested the effect of blocking Gβγ with the pharmacologic inhibitor gallein and by overexpression of Gα-transducin, a Gβγ scavenger (Crespo et al., 1994). Both treatments diminished pAktS473 and pERK1/2 levels in response to CRH (Fig. 5E,F), evidencing a role of Gβγ in CRHR1 signaling in this neuronal context.

We conclude that both SFK kinase family and the Gβγ complex participate in CRH/CRHR1 signaling. These results show that pharmacological or genetic inhibition of Src activity results in almost complete attenuation of CRHR1 signaling to phospho-AktS473. The higher degree of pERK1/2 inhibition with SFK inhibitors and the fact that only Akt phosphorylation was modulated by Src mutants may indicate that both activations could be mediated by different family members. Whereas pAkt levels seem to be dependent on Src activity, pERK1/2 could rely on another SFK kinase. Besides, the fact that Akt activation required Gβγ makes this complex a possible regulator of Src/SFK activation.

### CRHR1 endocytosis is required for Akt activation

The fact that in stimulated HT22-CRHR1 cells activated Akt profile overlaps the late phase of ERK1/2 activation, shown to be dependent on CRHR1 endocytosis and sAC activity (Bonfiglio et al., 2013; Inda et al., 2016), prompted us to examine if CRHR1 endocytosis and Akt activation may be linked. We tested this hypothesis using two experimental approaches to inhibit endocytosis: preincubation with Dyngo-4a, a pharmacologic inhibitor of dynamin GTPase activity and transient expression of a dominant-negative mutant of this GTPase (DynK44A). Notably, Dyngo-4a treatment completely blocked Akt phosphorylation (Fig. 6A). The reduced Akt activation caused by endocytosis inhibition was also evidenced by a diminished phosphorylation of Akt substrates (Fig. 6B). Transfection with DynK44A also reduced pAktS473 response compared with control cells (Fig. 6C).

**Figure 6:**
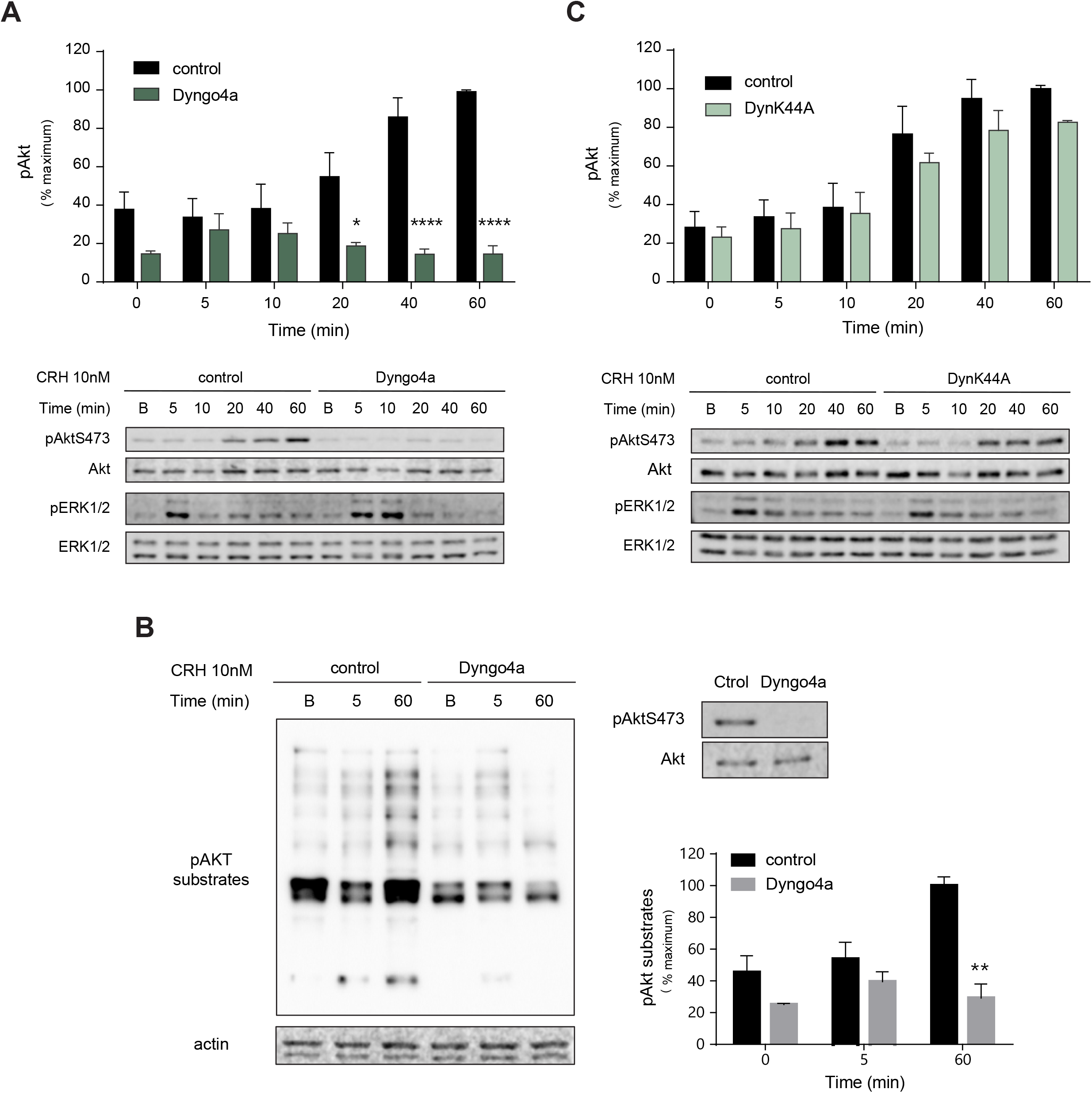
Akt activation by CRH depends on endocytosis. (A,B) HT22-CRHR1 cells were preincubated in the presence of vehicle (control) or 30μM Dyngo-4a. (C) Cells were transfected with dominant negative plasmid DynK44A. (A-C) Cells were stimulated with 10nM CRH for the indicated time points. Phosphorylated and total proteins were determined by immunoblotting. Signals were quantified by densitometry and each value of pAkt was normalized to total Akt. Results are expressed as percentage and relative to the mean of maximum pAkt in control condition (mean ± s.e.m, n=4). Two-way analysis of variance followed by Bonferroni’s test: *, p < 0.05; ****, p < 0.0001 with respect to control. Representative immunoblots are shown. B) pAkt substrates phosphorylation was immunodetected. Actin was detected as a loading control. pAktS473 and total Akt were detected as a control of inhibitor’s efficiency. Results are expressed as percentage and relative to the mean of maximum response in control condition (mean ± s.e.m, n = 3). Two-way analysis of variance followed by Bonferroni’s test: **, p < 0.01 with respect to control. Representative immunoblots are shown.

Recent work showed that chemical inhibitors of dynamin may exert additional inhibitory effects on signaling independent of the blockage of its target dynamin. These off-target functions were evidenced when genetic approaches to block dynamin gave opposite results (Basagiannis et al., 2017; Basagiannis et al., 2021; Park et al., 2013; Persaud et al., 2018). We have previously demonstrated that both Dyngo-4a treatment and DynK44A overexpression block CRHR1 endocytosis after ligand stimulation causing a diminished cAMP response and a blockage of the sustained ERK1/2 activation phase in HT22-CRHR1 cells (Bonfiglio et al., 2013; Inda et al., 2016). Although the different intensity of the effects of both treatments observed on Akt phosphorylation (Fig. 6A,C) may be explained by Dyngo-4a off-targets and/or by the efficiency of DynK44A expression, we conclude that in line with a late temporal profile of Akt phosphorylation by activated CRHR1, Akt activation is dependent on CRHR1 endocytosis.

Taking into account that activated CRHR1 internalization is fast, with a significant fraction of this receptor internalized after 5 min of CRH stimulation (Inda et al., 2016), endocytosis-dependent CRHR1 signaling mechanisms contribute to some extent to early activation events. In this scenario, the finding that both phases of ERK1/2 activation were diminished in the presence of PI3K/Akt inhibitors (Fig. 3C,D)-even before pAkt levels increased significantly in response to CRH- is consistent with Akt activation being dependent on CRHR1 endocytosis.

### Subcellular compartment localization of activated CRHR1

Identifying the nature of the subcellular compartments involved in CRHR1 trafficking is a starting point to understand the signaling cascades generated from locations different from the plasma membrane. To that end, we analysed the fluorescent signals of the N-terminal cMyc tag of CRHR1 in CRH-stimulated HT22-CRHR1 cells together with that of endogenously expressed proteins characteristic of endosomal compartments by immunofluorescence confocal microscopy. In unstimulated cells, CRHR1 mostly localized at the plasma membrane (Fig. 7, Basal) although some diffuse cytosolic staining was also detected. CRH stimulation resulted in receptor internalization evidenced as clusters of intracellular cMyc signal that were detected after 5 min, which became more conspicuous at 30 min of CRH treatment together with a decrease in plasma membrane signal (Fig. 7). We compared CRHR1 localization to that of three Rab proteins -Rab5, Rab11 and Rab7-used as markers of early, recycling and late endosomes respectively (Fig. 7A,B and Fig. S6A). Upon ligand stimulation, a proportion of Rab5 and Rab11 signal relocalized, forming aggregates that spatially overlapped CRHR1 clusters (Fig. 7A,B). This was not observed when assessing CRHR1/Rab7 (Fig. S6A). To quantify the degree of colocalization between fluorophores, the Manders overlap coefficient was calculated. In agreement with the qualitative analysis of the images, coefficient values increased with time in the case of Rab5/CRHR1 and Rab11/CRHR1, whereas only a very small fraction of internalized CRHR1 overlapped Rab7 endosomes (Fig. 7C, Fig. S6B).

**Figure 7:**
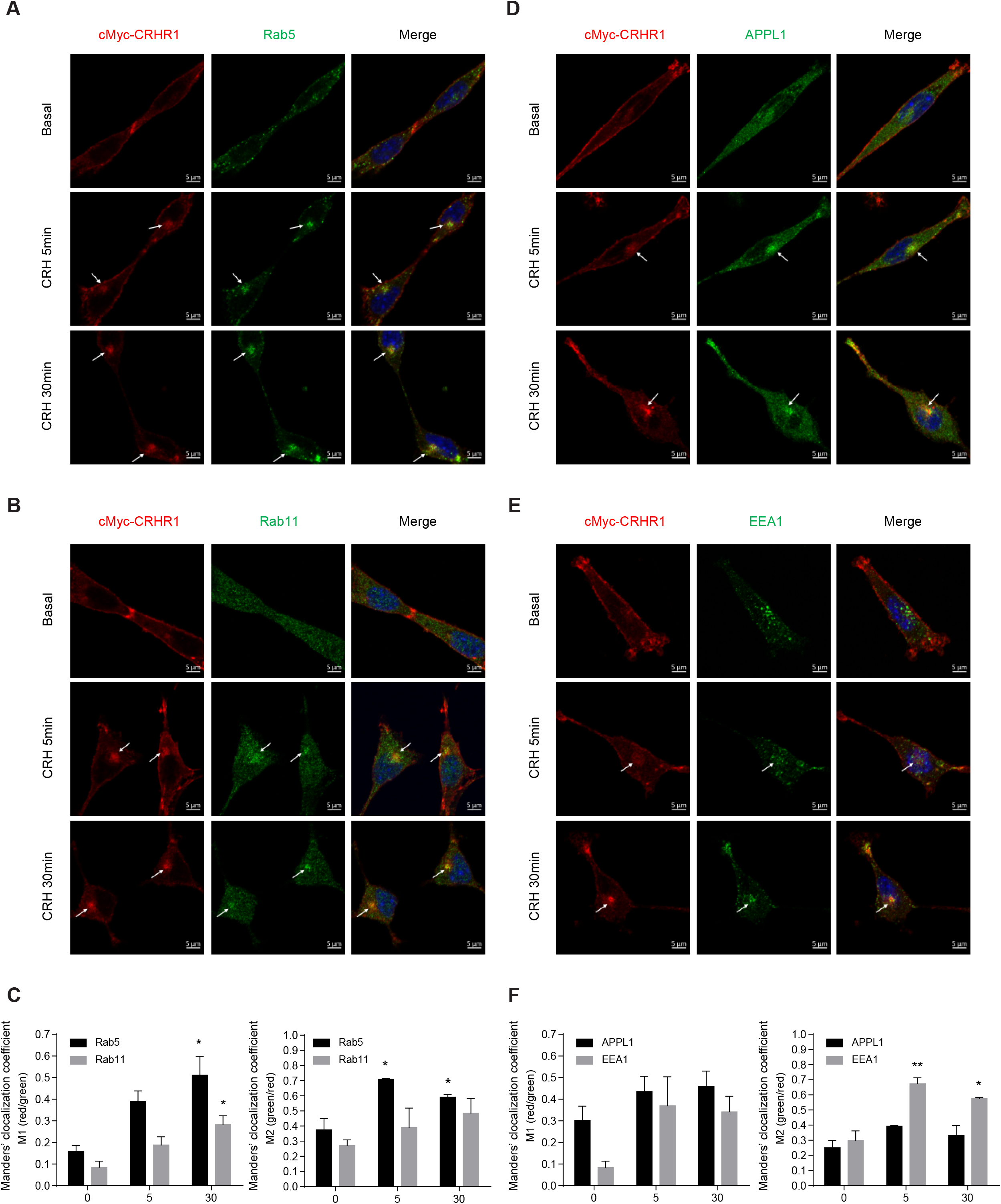
Activated CRHR1 colocalizes with early and recycling endosomal markers and Rab5 effectors. (A,B,D,E) HT22-CRHR1 cells were stimulated with 10 nM CRH for the indicated time points. cMyc-CRHR1 subcellular distribution was monitored together with Rab5 (A), Rab11 (B), APPL1 (D) and EEA1 (E) by indirect immunofluorescnce using specific antibodies. Representative confocal images of each condition are shown. Nuclei were stained with DAPI. Bars, 5µM. White arrows indicate internalized CRHR1 clusters. (C,F) Colocalization was measured quantitatively by Manders’ correlation coefficients (MCCs) M1 and M2 using ImageJ software with the JaCoP plug-in as described in Materials and Methods. M1 refers to the fraction of red channel signal (cMyc-CRHR1) overlapping green channel signal (Rab5, Rab11, EEA1 or APPL1) and M2 corresponds to the reciprocal fraction. Data: mean ± s.e.m, n = 3, 30-40 cells per condition. One-way analysis of variance followed by Tukey’s test: *, p < 0,05; **, p < 0,01; with respect to basal.

In the endocytic route, the complex molecular composition of early endosomes includes the downstream Rab5 effectors APPL1 and EEA1 that are involved in multiple intracellular signaling events (Sigismund et al., 2012). We investigated if internalized CRHR1 were transported to a specific compartment containing these early endosome characteristic proteins. In HT22-CRHR1 cells, CRHR1 immunofluorescence colocalized with both APPL1 and EEA1 signals of endogenously expressed proteins after 5 and 30 min of CRH stimulation (Fig. 7D,E). According to Manders coefficients values, the CRHR1 fraction associated to EEA1 notably increased after 5 min of CRH stimulation suggesting that the receptor seems to associate slightly more with EEA1 than APPL1 (Fig. 7F). Notably, silencing of APPL1 or EEA1 did not gave a noticeable change in CRHR1 intracellular distribution in CRH-stimulated cells (Fig. S7).

### A role for endosomal effectors APPL1 and EEA1 in CRHR1 signaling

Given that endocytosis is required for CRHR1-mediated sustained ERK1/2 activation (Bonfiglio 2013, Inda 2016) and for Akt signaling, and having observed that CRHR1 colocalized with both APPL1 and EEA1 proteins after CRH stimulation (Fig. 7D-F), we next explored if endosomal platforms containing these Rab5 effectors were involved in CRHR1 signaling from intracellular compartments in HT22-CRHR1 cells. To that end, we generated stable clones by transduction with non-target (control), APPL1 and EEA1 specific shRNA lentiviral particles. The effectiveness of the knockdown strategy of endogenous APPL1 and EEA1 in HT22-CRHR1 cells is shown in Fig. S8.

In HT22-CRHR1 cells, depletion of APPL1 with two different shRNAs (SH2, SH6) significantly attenuated CRH triggered Akt phosphorylation at S473 (Fig. 8B), consistent with a role of APPL1 in Akt signaling. In contrast, depletion of APPL1 did not impact on CRH-dependent ERK1/2 activation (Fig. 8A). Conversely, depletion of EEA1 with two shRNAs (SH4, SH5) diminished ERK1/2 phosphorylation downstream activated CRHR1 (Fig. 8C) without affecting Akt activation at Ser473 (Fig. 8D). These contrasting results indicate that depletion of each of these Rab5 effectors did not cause a general defect on signaling downstream CRHR1 but rather that APPL1 and EEA1 may regulate Akt and ERK1/2 signaling from discrete endosomal compartments in the neuronal context of CRHR1 activation.

**Figure 8:**
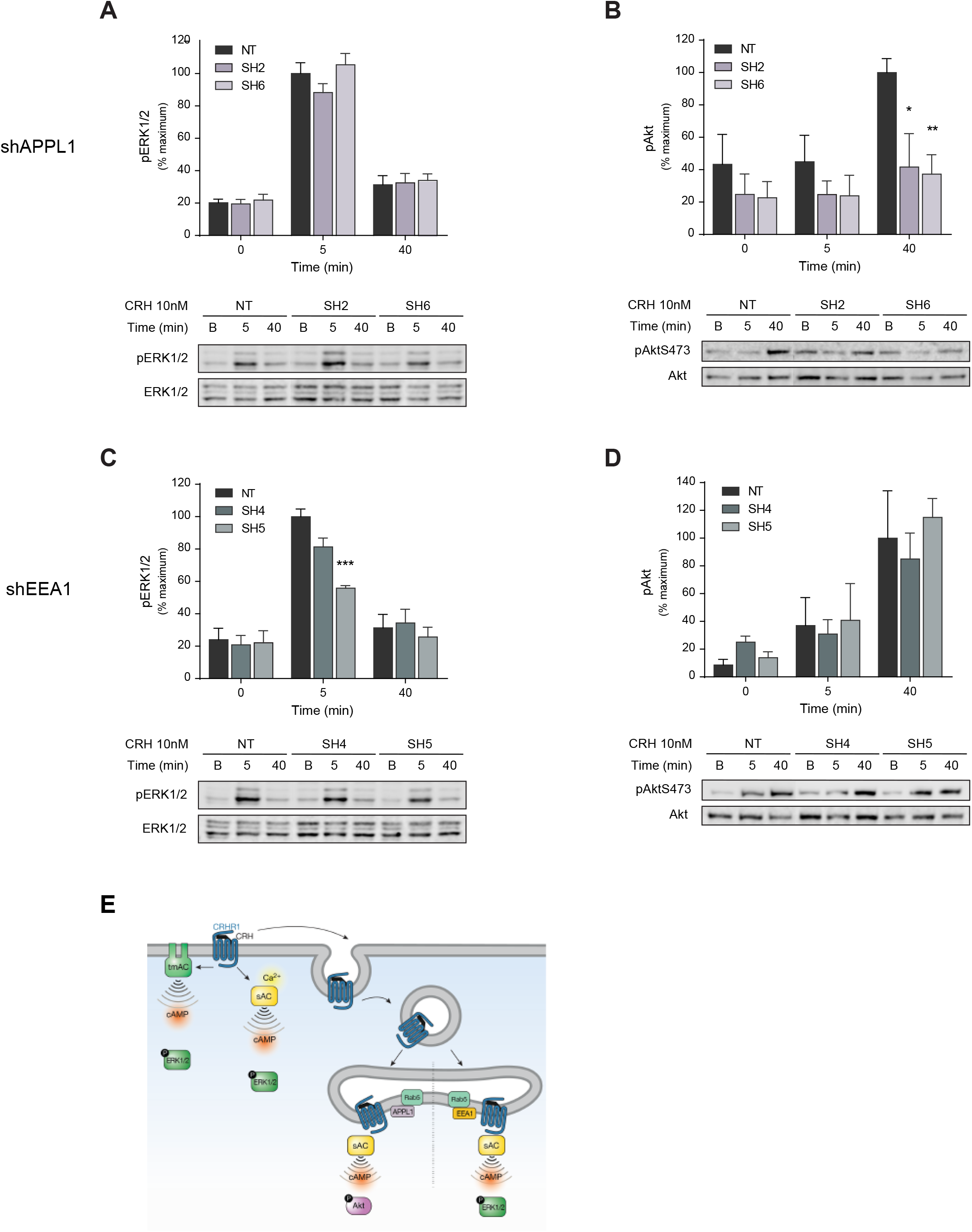
Effect of APPL1 and EEA1 silencing on CRHR1-mediated Akt and ERK1/2 activation. (A-D) HT22-CRHR1 cells stably expressing a control shRNA (NT) or APPL1-specific (SH2, 6) or EEA1-specific (SH4, 5) shRNAs were stimulated with 10 nM CRH for the indicated time points. Phosphorylated and total proteins were determined by immunoblotting. Signals were quantified by densitometry and each value of pERK1/2 (A, C), or pAkt (B, D) was normalized to total ERK1/2 or Akt, respectively. Results are expressed as percentage and relative to the mean of maximum response in NT condition (mean ± s.e.m, n = 3). Two-way analysis of variance followed by Bonferroni’s test: *, p < 0,05; **, p < 0,01; ***, p < 0,001; with respect to NT. Representative immunoblots are shown. E) Proposed model for CRHR1-dependent Akt and ERK1/2 activation. At the cell membrane, ligand stimulated CRHR1 triggers cAMP and intracellular calcium increase. tmACs and sAC activities contribute to ERK1/2 activation. Once internalized, CRHR1 continues to generate cAMP by the activity of sAC. Both sAC-generated cAMP pool and receptor internalization are essential for Akt and ERK1/2 activation from endosomes. In endosomal membranes, specific signaling complexes Rab5-APPL1 and Rab5-EEA1 are critical for Akt and ERK1/2 activation, respectively.

## DISCUSSION

In this study we characterized Akt activation downstream CRHR1 in hippocampal HT22-CRHR1 cells and its integration with molecular components and cellular mechanisms previously described by our work in this cellular context. We show that Akt is phosphorylated at Ser473 in response to CRHR1 high affinity ligands, CRH and UCN1, in a cAMP- and endocytosis-dependent manner.

The organization of cAMP signaling in micro/nano domains, in which the second messenger is selectively regulated at the subcellular level to ensure precise actions, contributes to the diversity and specificity of cAMP-regulated processes (Anton et al., 2022; Zaccolo et al., 2021). In HT22-CRHR1 cells, cAMP compartmentalization is achieved by the existence of two distinct cAMP pools: whereas both tmACs and sAC participate in cAMP production from activated CRHR1 at the plasma membrane, sAC is the key source of cAMP downstream internalized CRHR1 (Inda et al., 2016). In this work, we show that cAMP generated from CRHR1 activation through sAC is also essential for Akt activity, providing additional evidence of a functional diversification between tmACs and sAC (Pozdniakova and Ladilov, 2018; Valsecchi et al., 2017). To our knowledge, this is the first report on Akt activation downstream sAC activity, and one of the few linking sAC with GPCR-mediated kinase activation.

Akt activation downstream CRHRs stimulation has been poorly studied. Among the few reports available, most were performed in model cell lines with limited physiological relevance, and none explored cAMP involvement in the process probably because kinase activation was uncoupled from cAMP production in the cellular contexts studied. In most systems where CRHR1 functionality is crucial in pathophysiology, cAMP is a key mediator of CRH action (Inda et al., 2017a). In primary hippocampal and cortical neurons, we showed that CRH triggers a sustained cAMP response (Inda et al., 2016) which was similar to the one observed in stimulated HT22-CRHR1 cells, and different than the acute and transient response in AtT20 corticotrophs (Inda et al., 2016). Notably, cAMP production by CRHR1 stimulation in these cellular contexts is linked to MAPK activation (Inda et al., 2016). On the other hand, PI3K/Akt signaling associated to cAMP increase has been described for activated M1 muscarinic receptor in hippocampal neurons (Zhao et al., 2019) and hormone receptors such as FSHR, TSHR, LH and A2R (Alam et al., 2004; Cass et al., 1999; Gonzalez-Robayna et al., 2000; Phosri et al., 2017), although the spatial and temporal organization of the pathway is still not well understood. Recent work has also shown that phosphodiesterase 4 (PDE4) inhibition increased Akt activation in cortical neurons and HT22 cells exposed to oxygen deprivation (Xu et al., 2019), which is in line with cAMP-dependent Akt phosphorylation in neuronal contexts.

Interestingly, CRHR1 stimulation in HT22-CRHR1 cells activates ERK1/2 and Akt with opposed temporal profiles. While ERK1/2 activation occurs rapidly after stimulation, pAktS473 levels increase at time points that overlap the late phase of ERK1/2 activation, which depends on CRHR1 endocytosis and sAC activity (Bonfiglio et al., 2013; Inda et al., 2016). ERK1/2 and Akt activation by the CRH system has been reported in different cellular systems including non-neuronal cells (Blaabjerg et al., 2014; Chandras et al., 2009), AtT20 (Tsukamoto et al., 2010), COS7 (Parra-Mercado et al., 2019) and HEK293 cell lines (Punn et al., 2006). Akt and ERK1/2 response dynamics vary according to the cellular context analyzed, although activation profiles of both kinases were not analyzed in parallel. In this regard, the hippocampal context appeared as an interesting scenario in which the distinct phosphorylation kinetics of ERK1/2 and Akt suggested a possible crosstalk between their activities downstream CRHR1 in neurons.

We show that PI3K/Akt pathway modulates ERK1/2 activation in HT22-CRHR1 cells. Akt positive regulation of ERK1/2 activity has been evidenced downstream activated CRHRs overexpressed in COS7, HEK293 and CHO cells or endogenously expressed in AtT20, A7r5, CATHa cell lines (Brar et al., 2004; Markovic et al., 2011; Parra-Mercado et al., 2019; Punn et al., 2006; Tsukamoto et al., 2013). The opposite effect, MAPK ERK1/2 modulation of Akt activity, has not been deeply analyzed and showed different results depending on the CRHR and the cellular context studied. Regardless, these studies-like ours-provide evidence of the interplay between ERK1/2 and Akt activations downstream CRHRs. Although these pathways were initially considered as antagonistic, each regulating cellular programs that led the cell to opposite fates - proliferation vs differentiation, apoptosis vs survival – it has now been established that there is usually a great functional overlap between them. Cellular responses are therefore the product of their coordinated action, a fine balance between positive and negative cross-regulations.

Our results show that PI3K/Akt inhibition also affected the cAMP response and consequently CREB phosphorylation. cAMP levels can be modulated by modifying the activity of enzymes responsible for its production (adenylyl cyclases) or degradation (phosphodiesterases, PDEs). PI3Ks and Akt are classically considered cAMP negative regulators as they positively modulate PDE activity. For example, Akt phosphorylates PDE3A increasing its activity which lead to decreased cAMP levels and PKA activity (Han et al., 2006). PI3Kγ is a key regulator of β-Adrenergic receptor mediated contractility by regulating cAMP metabolism in the heart through PI3K kinase activity dependent and independent mechanisms. It is still unclear though, whether PI3Kγ promotes PDEs enzymatic activity or if it acts as a scaffolding protein for the assembly of multiprotein complexes that allow PDE recruitment to specific domains contributing to cAMP compartmentalization (Kerfant et al., 2006). In line with this, lymphocytic activation relies on an Akt/β-Arrestin/PDE4 complex recruited to lipid rafts by PI3K via Akt PH domain to mediate local cAMP degradation (Bjørgo et al., 2010).

Consistent with its temporal overlap with pERK1/2 late phase and its dependence on sAC-generated cAMP pool, our results reveal that CRHR1 mediated Akt activation and signaling require endocytosis. The dependence of Akt function on receptor internalization has been described for other GPCRs like the chemokine CXCR4 and PAC1R: interestingly, Akt mediated apoptotic effect in response to CXCL12 and PI3K/Akt neuroprotection in sympathetic neurons are affected by endocytosis blockage (English et al., 2018; May et al., 2010).

Spatiotemporal control is a usual mechanism that modulates activity and function of receptor signaling networks. This concept has changed our previous simplified view of cellular signaling from external stimuli as signaling emanating exclusively from activated receptors in the cell surface to a more complex model in which endocytosed receptors contribute to signaling cascades. In neuronal cells, stimulated CRHR1 exhibits prolonged signaling responses after internalization: cAMP originated from endosomal compartments by sAC accounts for a second sustained phase of ERK1/2 activation (Bonfiglio et al., 2013; Inda et al., 2016), and as we show here, is also required for Akt activity in HT22-CRHR1 cells reinforcing the functional association between sAC activity and endosome-based GPCR signaling.

Our studies show that Src and Gβγ are required for Akt activation by stimulated CRHR1 in HT22-CRHR1 cells, although the mechanisms involved remain to be determined. GPCR-mediated Src activation has been found to include RTK transactivation, direct GPCR-Src interactions or binding to GPCR associated proteins (Gα, Gβγ or β-arrestins). Our finding that Gβγ subunits can modulate pAkt levels makes of this complex a link between activated CRHR1 and Src in a hippocampal context. ERK1/2 and Akt activation via Src has been studied for ghrelin receptor and two mechanisms were described: one involving Gβγ and a second mediated by a β-arrestin associated complex (Camiña et al., 2007; Lodeiro et al., 2009). An intricate organization of β-arrestin protein complexes connect GPCR signaling to a variety of signaling pathways including ERK1/2 and Akt (Lefkowitz and Shenoy, 2005). Recently, β-arrestin-Src-GPCRs complexes have been demonstrated to allosterically activate Src (Pakharukova et al., 2020). Direct physical interaction between SFKs (Src and Fyn) and several GPCRs has been reported (Luttrell and Luttrell, 2004; McGarrigle and Huang, 2007). In HEK293 cells the 5-HT6R binds to Fyn, and ERK1/2 activation by receptor stimulation was found Fyn-dependent (Yun et al., 2007). Moreover, in COS7 cells, overexpressed CRHR1 and Src were detected in a complex with EGFR (Parra-Mercado et al., 2019). Future studies should address a possible physical interaction between CRHR1 and Src in a hippocampal context. Furthermore, given that CRHR1 prolonged signaling through ERK1/2 activation was found dependent on β-arrestin2 (Bonfiglio et al., 2013), further investigation of β-arrestin involvement in Akt activation will provide a connection between pathways activated by CRHR1 in neurons. Receptors traffic through a complex endocytic network composed of a variety of highly heterogeneous compartments, whose composition may vary according to context. Therefore, the protein repertoire available for signaling from these intracellular compartments dictates the spatiotemporal resolution of responses and determines receptor’s final fate within the network. A two-way relationship is established in which GPCR signaling and trafficking is regulated by the endocytic system which is in turn modulated by receptor-activated cascades. The link between CRHR1 signaling and trafficking reported in this work contributes to the identification of molecular components that define CRHR1 action from intracellular compartments.

Through immunofluorescence assays coupled with colocalization analysis we determined that CRHR1 traffics mainly to Rab5 early and Rab11 slow recycling endosomes in HT22-CRHR1 cells. Studies on CRHR1 trafficking in HEK293 cells detected CRHR1 in Rab5 endosomes, being receptor resensitization dependent on Rab11 activity, but not on Rab4 (Hasdemir et al., 2012). A previous report in the same cell line and in primary cortical neurons found that CRH-stimulated CRHR1 transits from Rab5 to Rab4 recycling endosomes (Holmes et al., 2006). CRHR1 trafficking characteristics may depend on the availability and abundance of molecular components involved in desensitization, internalization, and endosomal trafficking present in each cellular context analyzed. Although we have not ruled out Rab4 involvement in CRHR1 trafficking, our results are consistent with the existence of a slow recycling route for CRHR1 in the HT22 hippocampal neuronal context. Apart from indicating receptors trafficking routes inside the cell, Rab proteins can also determine specific signaling outcomes. Regarding PI3K/Akt pathway, a functional relationship has been established with Rab5 and Rab11 GTPases. Rab5 is required for insulin receptor 1 (IRS1)-p85α interaction that is a key step for PI3K/Akt activation (Su et al., 2006). PI3Kβ is a Rab5 effector (Christoforidis et al., 1999) whose interaction through p110β catalytic subunit is necessary for Rab5 to stay in its GTP-bound active state (Dou et al., 2013). On the other hand, Rab11 has been shown to interact in endosomes with the Gβγ complex in response to lysophosphatidic acid. This Rab11-Gβγ complex recruits PI3Kγ promoting Akt phosphorylation from intracellular compartments (García-Regalado et al., 2008). These data together with our immunocytochemical analysis not only support endosomal Akt activation but also place CRHR1 in intracellular compartments whose resident proteins can mediate this activation.

CRHR1 colocalized with both Rab5 effectors APPL1 and EEA1 in response to CRH. Experiments with knockdown cell lines presented alterations in receptor signaling: APPL1 depletion affected Akt activation with no effect on pERK1/2 levels, whereas EEA1 silencing had the opposite effect.

APPL1 is a multifunctional endosomal adaptor protein capable of promoting a variety of responses through interaction with different proteins. Relevant to this study are APPL1 interactions with Akt, PI3K, Rab5-GTP and receptors (Miaczynska et al., 2004; Mitsuuchi et al., 1999). In primary hippocampal pyramidal neurons Akt1 is transported retrogradely through axons in Rab5-APPL1 positive endosomes. APPL1 binds both Rab5-GTP and Akt1, this last dependent on endocytosis (Goto-Silva et al., 2019). APPL1-Akt endosomal interaction is also important in zebra fish embryo development. APPL1 loss induces widespread apoptosis in the embryo together with a decrease in pAkt and pGSK3β levels without affecting the phosphorylation state of other Akt substrates or the activation of MAPKs p38 and ERK1/2 (Schenck et al., 2008). These results show that APPL1 regulates not only Akt function but also its substrate specificity and hence the activation of associated cellular programs. Additionally, APPL1-mediated Akt activation is associated with a distinct intracellular location emphasizing the idea of signal compartmentalization by endocytosis.

Unlike APPL1, EEA1 is not a protein adaptor or scaffold but a PI3K regulated protein due to its ability to bind PI(3)P and a Rab5 effector necessary for membrane fusion (Simonsen et al., 1998). Membrane fusion requires Rab5-GTP, PI3K activity and Rab5-EEA1 interaction through EEA1 C-terminal end. Rab5 hyperactivity or structural changes in EEA1 C-terminal lead to abnormal endosomal structures (Patki et al., 1997; Simonsen et al., 1998). Interestingly, EEA1 has also been shown to dissociate from endosome membranes in response to PI3K inhibition by wortmannin (Patki et al., 1997). EEA1-positive endosomes have been also linked to GPCR-mediated Akt signaling: CXCR4 stimulation promotes endocytosis dependent activation of EEA1-Akt-FoxO3 pathway (English et al., 2018); Angiotensin II-mediated Akt activation in VSMCs relies on upstream kinases PKCα, PDK1 and p38 recruitment and activation in Rab-EEA1 endosomes (Nazarewicz et al., 2011). Recently, in neurons of the periaqueductal gray of chronic morphine-treated mice, phosphorylated ERK1/2 activated downstream β-arrestins was found to colocalize with EEA1 (Ram et al., 2021). In light of this evidence, the results obtained in this work could suggest that ERK1/2 intracellular activation may require either EEA1 action as an effector-like what was observed for APPL1 and Akt- or a structurally intact endosome.

Collectively, these results indicate that the endocytosis of ligand activated CRHR1 is critical for a complete cell response. At the membrane level, CRHR1 triggers cAMP rise from tmACs and sAC that leads to the activation of several cascades, i.e. pERK1/2. Once internalized, it continues to signal through cAMP, by the activity of sAC. Both sAC-generated cAMP pool and receptor internalization are essential for ERK1/2 and Akt activation at early and late time points after stimulation. The formation of specific signaling complexes in Rab5 early endosomes appears to be key for the cellular response of CRHR1. In fact, we identified different signaling platforms, being APPL1 critical for Akt activation and EEA1 for ERK1/2 activation, denoting the complexity of the endosomal network and signaling pathways (Fig.8E).

The sophisticated regulation of PI3K/Akt signaling is partly due to spatial and temporal compartmentalization including nanoscale membrane domains (Sugiyama et al., 2019). Thus, different Akt signaling nodes determine particular sets of Akt substrate regulation (Manning and Toker, 2017). Specific control of the wide variety of cellular responses elicited by Akt activation is crucial in pathophysiological processes. In the mature and developing brain, PI3K/Akt pathway regulates an impressive number of cellular functions, many linked to responses ascribed to the CRH system. Considering the potential interplay between the CRH system and Akt signaling, this work provides a basis for defining PI3K/Akt role in stress-related disorders and neuropathologies derived from CRH system dysregulation in the CNS.

## MATERIALS AND METHODS

### Cell culture and transfection

HT22-CRHR1 cells were cultured in DMEM supplemented with 5% FBS, 2 mM l-glutamine, 100 U/ml penicillin, and 100 μg/ml streptomycin (Invitrogen) at 37°C in a humidified atmosphere containing 5% CO2. Plasmids were transfected using Lipofectamine and Plus Reagent (Invitrogen) according to the manufacturer’s instructions and as previously described (Bonfiglio et al., 2013). Experiments were performed 48 h after plasmid transfection. Cells were co-transfected with a fluorescent protein expression vector to account for transfection efficiency. Only transfections yielding 60% efficiency or higher were used for further experiments.

### Plasmids and shRNA

Plasmids were provided as follows: the dynamin-K44A expression vector by J. Benovic (Thomas Jefferson University, Philadelphia, PA), N-EPAC by D.L. Altschuler (University of Pittsburgh School of Medicine, Pittsburgh, PA), Akt1-WT and Akt1-E17K by M. Blaustein (Instituto de Biociencias, Biotecnología y Biología, Buenos Aires, Argentina) and pCDNA3-Gα transducin, PKI-cherry, PKI mutant-cherry, pCEFL-SrcYF and pCEFL-SrcYF-KM by S. Gutkind (San Diego Moores Cancer Center, San Diego, CA).

shRNAs against APPL1 and EEA1 were selected from de Broad Institute GPP database and purchased from Sigma (MISSION TRC shRNAs purified plasmid DNA). A total of three individual shRNAs for each protein target were selected in plKO.1-puro backbone (SH1: catalog # SHCLND-NM_145221.2-220s21c1, TRC # TRCN0000340819; SH2: catalog # SHCLND-NM_145221.2-1890s21c1, TRC # TRCN0000340817; SH3: catalog # SHCLND-NM_001001932.3-1821s21c1, TRC # TRCN0000381960; SH4: catalog # SHCLND-NM_001001932.3-4866s21c1, TRC # TRCN0000381364; SH5: catalog # SHCLND-NM_001001932.3-349s21c1, TRC # TRCN0000256425; SH6: catalog # SHCLND-NM_145221.2-2339s21c1, TRC # TRCN0000340759).

### Generation of stable cell lines

HT22 stable clones expressing cMyc-CRHR1 (HT22-CRHR1) and FRET biosensors (HT22-cMyc-CRHR1-Epac-S^H187^ and HT22-CRHR1-AKAR4) were previously described (Bonfiglio et al., 2013; Inda et al., 2016).

Stable clones expressing both cMyc-CRHR1 and shRNAs against APPL1 or EEA1 were generated using Sigma MISSION shRNA lentiviral plasmids. HEK293T cells were cotransfected with plK0.1 shRNA vector and lentiviral helper plasmids (p.VSVG and p.D8.9) to obtain lentiviral particles. Particles were filtrated with 0.45µm pore size filter (Sartorius). HT22-CRHR1 cells were incubated 48 h with filtrated lentiviral particles of specific shRNAs for knockdown and a control (MISSION pLKO.1-puro Non-Target shRNA plasmid). Selection with Puromicin (3 μg/ml; InvivoGen) started 72 h post infection. After negative control (non-transfected HT22-CRHR1) died, clonal colonies were isolated and subcultured, and cells were maintained in culture with 1.5µg/ml puromycin. Knockdown efficiency was assessed by quantitative RT-PCR normalized to HPRT and by immunoblotting with specific antibodies (Fig. S7).

### Quantitative real-time PCR

Total RNA was extracted from cell lines using TRIzol reagent (Invitrogen), and complementary DNA synthesis was performed using M-MLV reverse transcription in the presence of RNasin RNase inhibitor (Promega). PCR primers were all intron spanning. cDNAs were amplified using Taq DNA polymerase (PBL). Quantitative real-time PCR was performed with SYBR Green I (Roche) using a CFX96 Touch Real-Time PCR Detection System (BIORAD). Relative expression was calculated for each gene by the Ct method with HPRT for normalization. Sequences and expected product sizes are as follows: *Eea1* forward 5’-CTTCAGACCAAAGCCCTAGAG-3’, reverse 5’-CGCCCACTTTCTGTTCAATG-3 (126 bp); *Appl1* forward 5’-CACCAAGTTTCCCATCAATCG-3’, reverse 5’-TCATTAAGTTTCCACCCTGCG-3’ (128 bp); *Hprt* forward 5’-TGGGCTTACCTCACTGCTTTC C-3’, reverse 5’-CCTGGTTCATCATCGCTAATCACG-3’ (139 bp).

### Ligand stimulation, drugs, and pharmacological inhibitors

When kinase activation (phosphorylation) was assayed, cells were serum-starved for 6 h in OptiMEM before drug pretreatments or stimulation. Cells were stimulated with human/rat CRH (4011473; Bachem), human UCNs (4027201, 4040984, 4039202; Bachem), PDGF (01-305 Millipore), forskolin (13019; Cayman Chemical), and 8-CPT-cAMP (F1221, Sigma-Aldrich) at the concentrations and time points indicated. After incubation, cells were washed with ice cold PBS and maintained in ice. When calcium chelators BAPTA-AM (B1205; Molecular Probes), EGTA (E3889; Sigma-Aldrich), or pharmacologic inhibitors were used, cells were pretreated with the drugs or vehicle 20 min before stimulation. The following inhibitors were used:, ddA (tmACs; D7408; Sigma), U0126 (MEK; 662005; EMD Millipore), H89 (PKA; 371962; Calbiochem), DIDS, anion channel blocker (D3514; Sigma-Aldrich), ESI09 (EPAC1/2; SML0814; Sigma), LY294002 (PI3K; 70920; Cayman), MK2206 (Akt; 11593; Cayman), gallein (Gβγ; 371708; Calbiochem), SU6656 (Src; S9692; Sigma), Dasatinib (SFKs; CDS023389; Sigma), Saracatinib (SFKs; AZD0530; Selleck Chemicals) and Dyngo-4a (dynamin inhibitor; ab120689; Abcam). For experiments testing bicarbonate dependence, bicarbonate-free DMEM supplemented with 25 mM Hepes was used to prepare media at 0- and 25-mM sodium bicarbonate concentrations. Each condition was adjusted to physiological pH 7.2 with NaOH and HCl and allowed to equilibrate in the incubator for additional 30 min, and then the pH was checked again and adjusted to 7.4. HT22-CRHR1 cells were serum-starved for 6 h in bicarbonate-free DMEM or 25 mM bicarbonate DMEM before stimulation with 10 nM CRH.

### Preparation of cellular extracts and immunoblotting

After treatments, cells were washed with ice-cold PBS and lysed in Laemmli sample buffer. Whole-cell lysates were sonicated and heated to 95°C for 5 min. Samples were resolved by SDS-PAGE and transferred to 0.45-nm nitrocellulose membranes (GE Healthcare Lifescience, Germany) for immunoblotting. Membranes were blocked in TBS–Tween 20 (0.05%) containing 5% milk at RT for 1 h under shaking and probed overnight at 4°C with the primary antibodies. The following antibodies were used: anti–pERK1/2 (E-4; sc-7383) and anti– β-actin (C4; sc-47778) from Santa Cruz Biotechnology, Inc.; anti–total-ERK1/2 (9102), anti-pAkt473 (4058), anti-total-Akt (2920), anti-α-tubulin (2144), anti-EEA1 (3288), anti-APPL1 (3858), anti-pAkt-substrates (9614), anti-pGSK3 (9331), anti-pS6 (2215), anti-S6 (2317) from Cell Signaling Technology, anti-phospho CREB (06-519; EMD Millipore).

Chemiluminescent signals were detected by HRP-conjugated secondary antibodies (BIORAD) and enhanced chemiluminescence (SuperSignal West Dura; Thermo Fisher Scientific) using a GBOX Chemi XT4 (Syngene). Fluorescent signals were detected with the Odyssey Fc Imaging System (LI-COR Biosciences), using anti-mouse-IRDye680LT, anti-mouse-IRDye800CW, anti-rabbit-IRDye680LT and anti-rabbit-IRDye800CW secondary antibodies (LI-COR Biosciences). Phosphorylated proteins were relativized to the total protein level, and results were expressed as the percentage of maximum response after stimulation. Immunoreactive signals were analyzed digitally using ImageJ software (National Institutes of Health).

### Spectral FRET

HT22-CRHR1 cells stably expressing FRET biosensors were seeded in glass-bottom dishes. Cell imaging was performed on an inverted LSM 710 (ZEISS) equipped with an automated stage, an incubation chamber, and Definite focus system. Data acquisition was performed with ZEN Black 2011 software. Images were acquired with a C-Apochromat 40×/1.2 water immersion and temperature-corrected objective lens at 1,024 × 1,024 pixels, 16 bits, pixel dwell time 3.15 μs, with open pinhole (600 μm).

The emission spectra of the fluorescent proteins Cerulean, mTurquoise2 and Venus were determined using 405 or 458 nm lasers for the excitation and lambda mode for the detection using a 32-channel QUASAR detector arranged with bandwidth channels of 9.7 nm.

For FRET experiments, cells were illuminated with a 30 mW, 405 nm diode laser at 2% laser power (550–650 gain) or a 25 mW, 458 nm argon laser at 3% power (650-750 gain) and a 405-nm or 458 nm dichroic mirror, respectively; emission was collected at 413 to 723 nm wavelengths every 15 s. The saturation level was verified for each image. Due to the intrinsic fluorescence of MK-2206, experiments using this inhibitor were performed with the 458 nm laser.

For the reference spectra and the experiments, phenol red–free DMEM/F12 medium supplemented with 20 mM Hepes was used, and imaging was performed at 37°C and 5% CO2. Approximately 2.5 min after the start of the experiment, 10 nM CRH was added. When indicated, cells were preincubated for 20 min with different reagents. Inhibitors were also added after CRH stimulation to evaluate the change in the FRET time course. For lambda FRET experiments, the linear unmixing tool of ZEN Black 2011 software was used to obtain an image for each fluorophore using the reference spectra. Image quantification was performed with Fiji software. After background subtraction, FRET and donor intensities were measured for single cells for each time point. The FRET/donor ratio was calculated and normalized to resting levels, before stimulation (dos Santos Claro et al., 2019).

### Immunofluorescence and confocal microscopy

HT22-CRHR1 cells were seeded on glass coverslips. After treatment, cells were washed with ice-cold PBS (supplemented with Ca^+2^ and Mg^+2^), fixed with 4% paraformaldehyde in PBS, and permeabilized with 0.01% Triton X-100/PBS. After washing with PBS, nonspecific binding was reduced by incubating coverslips with 5% FCS in PBS for 1 hour. Primary antibodies were diluted (1:150, cMyc; 1:100, pAktS473, EEA1, Rab5, Rab7; 1:50, Akt, APPL1, Rab11) and incubated for 2h in a humidity chamber at room temperature. After washing with 0.05% Tween/PBS, cells were incubated with a 1:200 dilution of Alexa Fluor 647 goat anti-mouse (A21235) or Alexa Fluor 488 donkey anti-rabbit (A21206) secondary antibodies (Molecular Probes) for 1h in a humidity chamber at room temperature. Finally, cells were washed with 0.05% Tween/PBS, stained with DAPI, and mounted with Mowiol on microscope slides. When double staining was carried out, both primary and secondary antibodies were incubated together. Autofluorescence, controls without primary antibody and single staining for bleed through controls were performed for each experiment.

Images were acquired on an inverted LSM 710 (ZEISS). Data acquisition was performed with ZEN Black 2011 software. Images were acquired with a C-Apochromat 40×/1.4 DIC M27 oil immersion objective lens at 984 × 984 pixels, 8 bits, pixel dwell time 3.29 μs, with 1 Airy unit. For each treatment, 20–40 individual cells were randomly selected and examined. No fluorescence was observed in cells treated with secondary Alexa-Fluor antibodies only or with no antibodies (autofluorescence). All images were acquired with the same confocal settings to ensure uniformity detection for each cell, including pinhole size, detector gain, offset, and laser power. The following antibodies were used: anti–cMyc (9E10; sc-40) from Santa Cruz Biotechnology, Inc.; anti-EEA1 (3288), anti-APPL1 (3858), anti-Rab5A (46449), anti-Rab11 (5589), anti-Rab7 (9367) from Cell Signaling Technology.

### Image analysis

Image processing and analysis were performed using the open source software FIJI and CellProfiler. Cell segmentation was performed with CellProfiler. First, color images were split into three channels (Akt, pAkt and DAPI) and transformed into 8-bit using FIJI given that CellProfiler works on grayscale images only. A predesigned pipeline from CellProfiler was modified to perform the desired analysis. Nuclei segmentation was performed with the module “IdentifyPrimaryObjects” on DAPI images while entire cells were segmented with the module “IdentifySecondaryObjects” on the Akt or pAkt images. For each object, integrated fluorescent intensity was measured and each value of pAkt signal was normalized by the corresponding total Akt to correct for Akt expression levels. Colocalization of cMyc-CRHR1 with different endocytic compartment markers was determined calculating the Manders correlation coefficients (M1 and M2) with the FIJI JACoP plugin. Images fed into de plugin must contain one cell per field and good signal intensity in the two channels being analyzed. Once the images are loaded, “M1 & M2 coefficients” is selected from the analysis list and thresholds for each image are set manually before performing the analysis.

### Statistics

Each experiment was performed at least three independent times. The results are presented as the mean ± s.e.m of each measurement. Comparisons between treatments were performed using one- or two-way analysis of variance (GraphPad Prism) followed by Bonferroni’s or Tukey’s test. Statistically significant differences are indicated.

## ACKNOWLEGMENTS

We thank E. Arzt for generous scientific support, V. Burghi and S. Gutkind for contributing constructs, R. M. Biondi and M. Blaustein for contributing reagents, and providing critical advice for experimental development and data interpretation.

## AUTHOR CONTRIBUTIONS

Conceptualization: P.A.dSC., S.S; Methodology: P.A.dSC., A.A., C.I.; Software: P.A.dSC., A.A., C.I. Validation P.A.dSC., N.A., K.E.L., M.S., Formal analysis: P.A.dSC., N.A., A.A; Investigation: P.A.dSC., N.A.; Resources: P.A.dSC., N.A.; Data curation: P.A.dSC., A.A.; Writing - original draft: P.A.dSC., S.S.; Writing - review & editing: P.A.dSC., C.I., N.A, S.S., Visualization: P.A.dSC., C.I., S.S. Supervision: S.S.; Project administration: S.S.; Funding acquisition: S.S.

## Supplementary information

Supplementary information available on line as

